# UMI-Gen: a UMI-based reads simulator for variant calling evaluation in paired-end sequencing NGS libraries

**DOI:** 10.1101/2020.04.22.027532

**Authors:** Vincent Sater, Pierre-Julien Viailly, Thierry Lecroq, Philippe Ruminy, Caroline Bérard, Élise Prieur-Gaston, Fabrice Jardin

## Abstract

**Motivation:** With Next Generation Sequencing becoming more affordable every year, NGS technologies asserted themselves as the fastest and most reliable way to detect Single Nucleotide Variants (SNV) and Copy Number Variations (CNV) in cancer patients. These technologies can be used to sequence DNA at very high depths thus allowing to detect abnormalities in tumor cells with very low frequencies. A lot of different variant callers are publicly available and usually do a good job at calling out variants. However, when frequencies begin to drop under 1%, the specificity of these tools suffers greatly as true variants at very low frequencies can be easily confused with sequencing or PCR artifacts. The recent use of Unique Molecular Identifiers (UMI) in NGS experiments offered a way to accurately separate true variants from artifacts. UMI-based variant callers are slowly replacing raw-reads based variant callers as the standard method for an accurate detection of variants at very low frequencies. However, benchmarking done in the tools publication are usually realized on real biological data in which real variants are not known, making it difficult to assess their accuracy.

**Results:** We present UMI-Gen, a UMI-based reads simulator for targeted sequencing paired-end data. UMI-Gen generates reference reads covering the targeted regions at a user customizable depth. After that, using a number of control files, it estimates the background error rate at each position and then modifies the generated reads to mimic real biological data. Finally, it will insert real variants in the reads from a list provided by the user.

**Availability:** The entire pipeline is available at https://gitlab.com/vincent-sater/umigen-master under MIT license.

## 1 Introduction

Old sequencing technologies are quickly disappearing as their use is relatively expensive, slow and doesn’t allow sequencing at very high depths. Due to their limits, they are now replaced by Next Generation Sequencers such as Thermo Fisher or Illumina. Prior to sequencing, DNA must be extracted and amplified by PCR in order to generate enough fragments to cover the wanted amplicons. After amplification, the sequencer handles the obtained fragments and generates their sequences in form of reads. The obtained reads must then be aligned to a reference genome in order to be used effectively. Today, cancer diagnosis is a very active area of research and one of its most important applications is the detection of Single Nucleotide Variants (SNV) in tumor cells. In fact, each cancer type has a specific profile of genetic mutations in specific genes. Therefore, establishing a precise profile of variants in a cancer patient allows us to better understand the cancer evolution and let us customize the treatment according to the established profile.

Performing variant calling analysis on the aligned reads is the step in which variants are called. Generally, variant calling tools can detect mutational events such as substitutions, insertions and deletions very efficiently. However, at very low frequencies (under 1%), it becomes very challenging for raw-reads-based variant callers to accurately call variants. In fact, PCR amplification and the sequencing step can introduce errors in the final reads. These errors are called artifacts and occur at very low frequencies which can lead to the confusion between them and true low-frequency variants. Multiple studies (Schmitt *et al*. (2012), Kukita *et al*. (2015), Newman *et al*. (2016), Young *et al*. (2016) and Bar *et al*. (2017)) have shown the effectiveness of using Unique Molecular Identifiers as a way to filter out PCR and sequencing artifacts. UMIs are short arbitrary oligonucleotide sequences that are attached to DNA fragments by ligation before the PCR amplification. The UMI tags must be arbitrary sequences so each fragment can have a unique short oligonucleotide sequence attached to it, giving each fragment a unique sequence tag. During the amplification, the UMI tags are amplified with their respective fragments. After sequencing, each unique UMI tag can be figured out from the reads. The idea behind using UMI tags in NGS experiments to filter out artifacts is explained in Figure 1. In fact, if a variant is a true mutation, it means that it must have been present on the initial DNA fragment so when we tag the DNA fragment with a UMI, we are also tagging the mutation. The fragments that result from the amplification of that mutated DNA fragment must all be tagged by the same UMI tag and carry the same mutation (Figure 1A). On the other hand, if the variant is a sequencing error, it means that the initial DNA fragment didn’t have the mutation in the first place and that it appeared later in the sequencing step. Therefore, during the amplification step, all the fragments resulting from the amplification of that DNA fragment should be tagged with the same UMI and should not present the mutation. The mutation will be produced later on, in the sequencing step, affecting only some reads and not all of them, thus creating discrepancies in the same UMI group (Figure 1B).

**Fig. 1.**
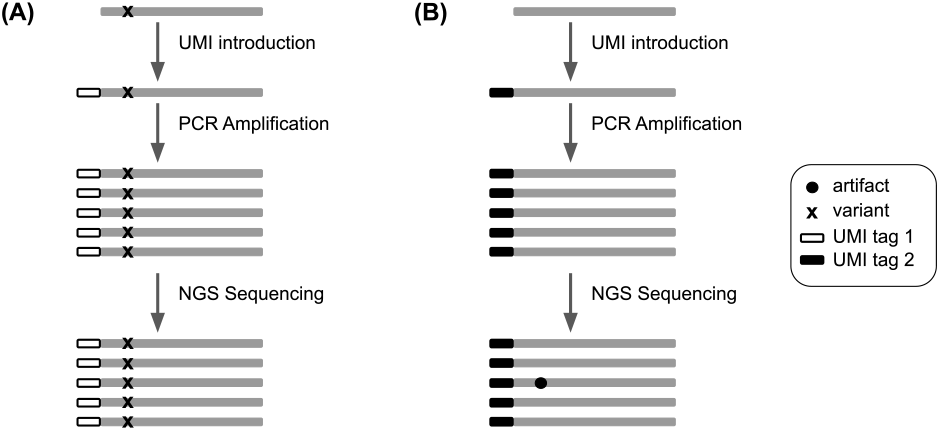
The difference between a true variant and an artifact from a UMI perspective. (A) A true variant is present on the DNA fragment so when the UMI tag 1 is added, it tags the fragment and the mutation as well. After amplification, all the fragments tagged with the UMI tag 1 carry the same mutation. (B) An artifact is not present on the DNA fragment but rather appears at the steps that follow the UMI introduction. That’s why not all fragments with the same UMI tag 2 carry the same artifact.

With the growing number of variant calling tools, it has become hard to choose the right tool adapted to a certain experiment. Data simulation can play an important role for testing different tools on a dataset that we have control on, a control that we don’t have on real biological data. Many short reads simulators exist at the moment. These tools are publicly available for researchers and allow them to test their algorithms on a simulated dataset, a dataset in which variants are inserted at different frequencies and at different positions. The usage of the reads simulators enable having a very accurate benchmarking of each variant calling tool ability. Surprisingly, no simulation software exists at the moment that let users generate reads with UMI tags. In this article, we present UMI-Gen, a UMI-based read simulator that can be used not only to test raw-reads based variant callers but most importantly, UMI-based ones. UMI-Gen uses multiple real biological samples to estimate background error rate and base quality scores at each position. Then, it will introduce real variants in the final reads. To test our tool, we used 6 control samples and show exactly how our algorithm estimate the background error rate at each position. Then we give it a list of 15 variants at different positions and at different frequencies to introduce them in the final reads. Finally, we used 2 raw-reads-based variant callers: SiNVICT (Kockan *et al*. (2017)) and OutLyzer (Muller *et al*. (2016)) and two UMI-based variant callers: DeepSNVMiner (Andrews *et al*. (2016)) and UMI-VarCal (Sater *et al*. (2020)) in order to compare the 4 tools performance and demonstrate that UMI-Gen correctly inserts the given variants at their respective positions and at the correct frequencies.

## 2 Materials and methods

### 2.1 Software input

UMI-Gen requires a minimum of three parameters at execution: a list of control BAM/SAM samples, the BED file with the coordinates of the targeted genomic regions and a reference genome FASTA file with BWA index files. UMI-Gen can also accept a fourth optional file under the PILEUP format. In fact, when running UMI-Gen on control samples, a PILEUP file is automatically produced. This file contains the A, C, G and T average counts at each position for all the control samples. This file can be given to UMI-Gen at execution time and will allow the software to reload the pileup generated during the last analysis instead of regenerating it. This will allow the user to gain some significant time since the pileup generation is the most time-consuming step.

#### 2.1.1 Control samples

Control samples are BAM/SAM files that are obtained by sequencing healthy individuals and normally shouldn’t contain any somatic variant. UMI-Gen can accept input files in BAM and SAM formats. A pileup is performed on each sample and a final average pileup is generated from the counts of all control samples.

#### 2.1.2 Variants file

This file contains a list of the variants the user wishes to insert in the simulated reads. These are the only variants that should be reported in the variant callers VCF file during variant calling benchmarks. The variants file is a Comma Separated Values (CSV) file that contains 2 columns: the first column contains the variant ID with the HGVS nomenclature and the second column being the variant’s desired frequency. UMI-Gen will then go to each position and insert these variants in order to produce final reads.

### 2.2 Generating the final pileup

#### 2.2.1 Pileup

The first step of the workflow (Figure 2) consists of generating the *final_pileup*. For each control sample, our pileup algorithm will count the occurrences of each A, C, G and T. The counts will be stored for each position of the BED file as well the average quality of the position and its depth. This is basically the same algorithm that is used by UMI-based variant caller UMI-VarCal that’s been reintegrated in this tool for its high efficiency in treating reads with UMI tags. When all the pileups for all the control samples are ready, they will be merged in a final pileup that contains the average statistics (counts, depth and quality score) at each position based on the observations on all control samples (Figure 2A). When the *average_pileup* is complete and ready, it will be automatically dumped as a PILEUP file that contains all the calculated information on the set of controlled samples. If the user wishes to generate simulated data based on the same BED file and the same set of control samples, the dumped pileup can be used directly which allows the program to skip the pileup generation step and go directly to the variant calling step, saving the user much significant time.

**Fig. 2.**
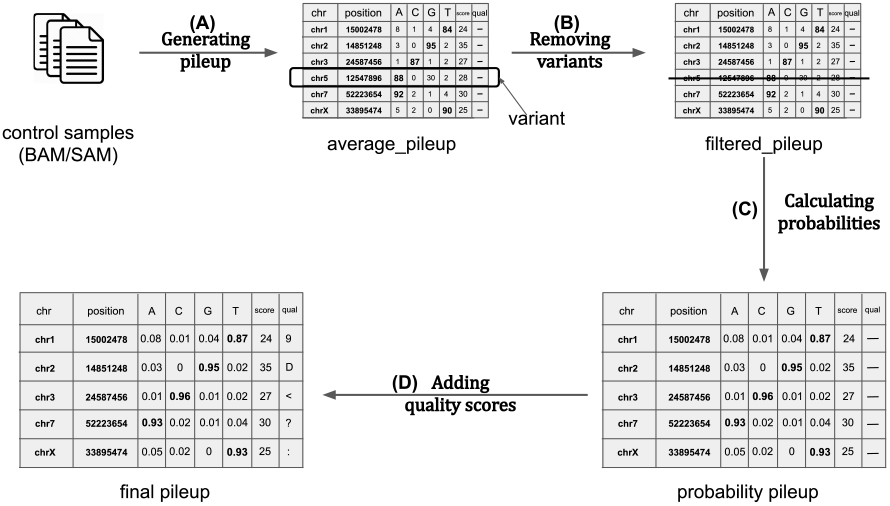
Background error estimation workflow. (A) The first step runs over every position in all control samples and counts the total occurrences of every A, C, G and T. It also stores the average base quality score for each position. (B) The second step’s goal is to remove any suspected variant from the pileup as our objective is to estimate background error noise only. (C) In this step, the counts are converted to probabilities by dividing them by the depth for each position. (D) The final step consists of converting the base quality score of each position to the corresponding ASCII + 33 character.

#### 2.2.2 Variant Calling

Even though the control samples are theoretically variant-free, SNP and undetected mutations could still be present in the files. These potential variants must be removed so they would not be present in the final reads. To do so, we used the same variant calling method implemented in UMI-VarCal to call out potential variants and remove them from the pileup. This step will produce what we call a *filtered_pileup* (Figure 2B).

#### 2.2.3 Background noise estimation

The background noise estimation step consists of calculating the frequency of observing an A/C/G/T at each position. Without the background errors, at each position the reference base should have a frequency of 1 while remaining three bases should be at 0. The total of the four frequencies must be equal to 1. However, we know that artifacts exist in our control samples and these artifacts represents the background noise that we normally encounter in a normal NGS experiment. Since our aim is to simulate reads that resemble the most to reads produced under real and normal sequencing experiments, UMI-Gen calculates the real base frequencies from the control samples at each position. The frequencies will then be used as a probability matrix when producing the final reads. When this step is complete, a *probability_pileup* is generated (Figure 2C). Insertions and deletions are not considered during the background noise estimation and thus, are not present in the *final_pileup* as their occurrence is very low especially in second and third generation sequencers. Therefore, we judge that their inclusion is not worth complicating the algorithm for.

#### 2.2.4 Quality scores estimation

Our tool was developed on sequencing files produced by an Illumina sequencer. In the FASTQ files produced by Illumina sequencers, quality scores are encoded into a compact form, which uses only 1 byte per quality value. In this encoding, the quality score is represented as the character with an ASCII code equal to its value + 33. The lowest quality score of a base can be 0, the corresponding ASCII code is 33 and the ASCII character that is found in the FASTQ quality string is ‘!’. The highest quality score is 41 and the corresponding ASCII character is the letter ‘J’. The full table of correspondence is available in Table S1. UMI-Gen is therefore only compatible with sequencers that use the same encoding. UMI-Gen calculates the average quality score for each position based on the qualities in all control samples and then converts the quality score to the corresponding ASCII character to be inserted in the final FASTQ file. This is the final step of the pileup generation workflow and will produce the *final_pileup* (Figure 2D).

### 2.3 Producing the reads

The main objective of UMI-Gen is to generate paired-end reads that mimic reads obtained from real life experiments. That’s why, it starts exactly the way a real-life sequencing experiment starts: getting the DNA fragments. At the beginning, our tool will generate a number of initial sequences that only present the reference base at each position. Then, a unique UMI tag is attached to each initial sequence. Depending on the amplification factor and the desired depth chosen by the user, the algorithm will keep amplifying the initial sequences until the desired depth is reached at all positions. Once we have the reference reads, the second step consists of adding the background noise (refer to section 2.2.3) to these reads (Figure 3A). Using the probability matrix calculated before, UMI-Gen modifies the reads at each position for them to match the calculated probabilities. These modifications are done without changing the reads UMI so they mimic PCR and sequencing artifacts: they are false positives and should not be called by variant callers. Finally, UMI-Gen parses the variants file provided by the user in order to insert true mutations in the final reads. The algorithm will go to each position, change the probability of the variant to the corresponding frequency from the variants file. In this step, since UMI-Gen is adding a true variant, the UMI tags of the modified reads are also modified in order to produce concordant UMI tags (Figure 3B). A concordant UMI tag is a UMI whose all reads carry the exact same mutation. Also, since UMI-Gen generates paired-end data, when adding a mutation on one read, the variant is automatically added to its mate.

**Fig. 3.**
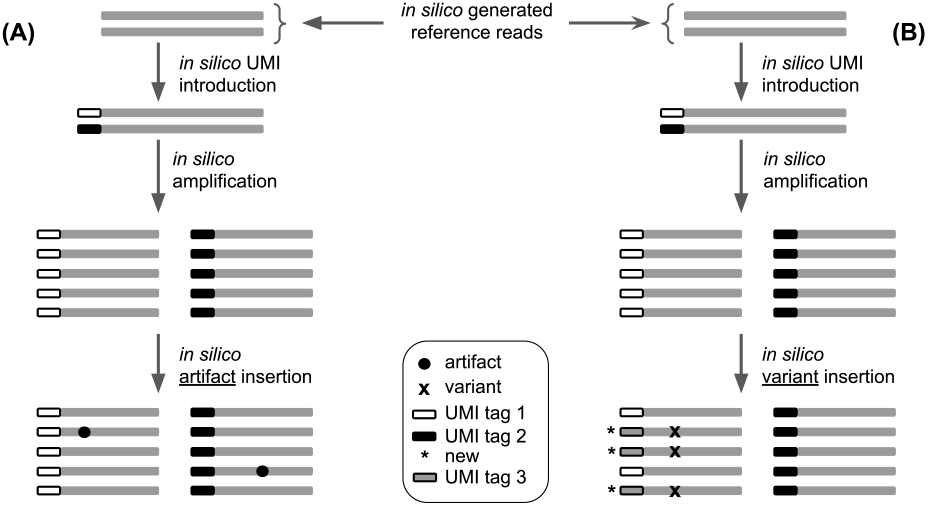
The difference between adding a true variant and adding an artifact in generated reads. (A) Adding an artifact is relatively easy as all the tool has to do is to modify the base at the wanted position without touching the read’s UMI tag. (B) On the other hand, in order to add a true variant, the software must change the base at the wanted position on a set of reads. Then it will create a new UMI tag (UMI tag 3) and change the UMI tag of all the affected reads to UMI tag 3.

### 2.4 Software output

Once all variants are inserted, UMI-Gen will generate the two FASTQ files (R1 and R2). It will then call BWA (Li and Durbin (2009)) to do the alignment, a step that will produce a BAM file. SAMtools (Li *et al*. (2009)) is finally called to create the BAM’s index file and convert the BAM into SAM. All five files are generated in the desired output directory. In addition, UMI-Gen generates a binary PILEUP file that corresponds to the average pileup dumped. This file can be used to skip the pileup regeneration and load the pileup directly if the analysis was already done on the same control samples.

### 2.5 Implementation

Launching UMI-Gen’s workflow (Figure 4) is handled by a main Python script that controls many Python3 modules. In order to achieve better overall performance, Cython was used to compile all Python modules. UMI-Gen requires for the tools BWA and SAMtools to be installed on the PC/server: BWA is called for the alignment step and SAMtools for converting, sorting and indexing the generated BAM files. Our tool can be executed through a UNIX/Linux command line interface. In total, UMI-Gen can accept 20 parameters at execution. Managing these parameters allows the user to have full control over his simulated data. A list of all the parameters and thresholds is available in Table S2.

**Fig. 4.**
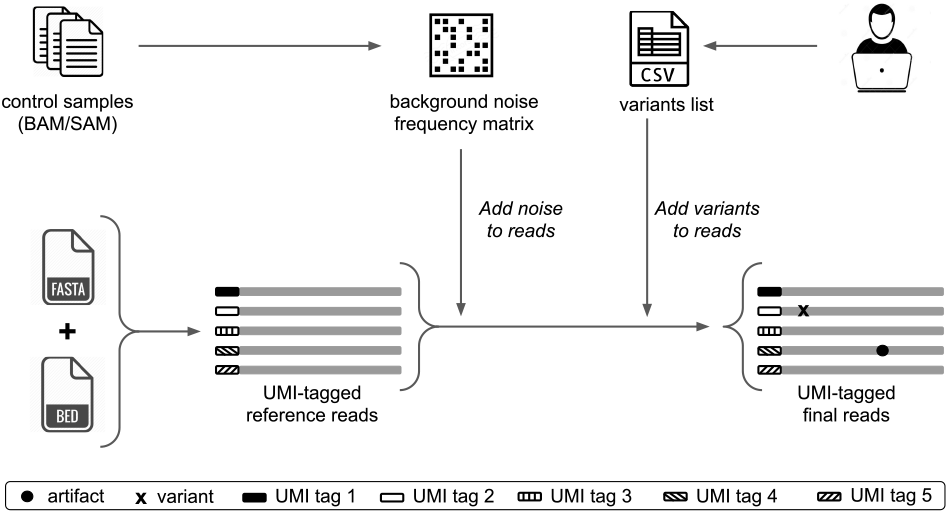
UMI-Gen’s workflow: Control samples are used to create a background noise frequency matrix and the user provides a CSV file with a list of the wanted variants. Using the FASTA and the BED files, UMI-Gen creates a first set of UMI-tagged reference reads. Artifacts are then inserted to mimic the sequencer’s background noise. Finally, the tool uses the list provided by the user to insert variants at their exact locations.

## 3 Results

### 3.1 Control samples

A targeted sequencing panel was designed at the Centre Henri Becquerel in Rouen (France) to search for specific mutations in the DNA of patients suffering from Diffuse Large B cell Lymphoma (DLBCL). This panel of 76 630 bases is designed to identify genomic abnormalities within a list of 36 genes that are most commonly impacted in this type of lymphoma. The panel was specifically designed for QIAseq chemistry allowing UMI introduction in the DNA fragments during the construction of the library. A list of the genes used in the panel and their corresponding number of targeted regions is provided in the supplementary Table S3. In order to test our tool’s ability to mimic and reproduce average sequencer background noise in the produced sample, we randomly selected 6 samples from a very large number of patients whose DNA were sequenced at the Centre Henri Becquerel. All six samples are liquid biopsies with circulating cell-free DNA that was checked to be adequate for sequencing. We preferred the use of liquid biopsies as these samples tend to be loaded with very low frequency variants and artifacts. Using such samples as control samples will produce simulated data with a relatively high number of artifacts. This will allow us to have an accurate estimate of the specificity of each tested variant caller.

To make sure that the artifacts were correctly added to the reads, we used IGV (version 2.4.16) (Robinson *et al*. (2011)) to visualize the reads. Figure 5 shows the exact counts of A, C, G, T for the position 2 493 165 on the chromosome 1 for each control sample. The fist control sample counts (0,11,10,874), the second sample has (0,1,7,843), the third one has (0,2,2,860), the fourth sample shows (1,6,9,965), the fifth one has (1,2,4,867) and the final one counts (3,2,2,880). As explained in section 2.2.3, UMI-Gen will calculate an average count for each base and then estimate its probability. In our case and for this position, the obtained average count has 4 A, 24 C, 34 G and 5289 T with a total count of 5351 bases. To obtain the probabilities for this position on this chromosome, we simply divide each base count by the total count of the 4 bases, obtaining the final probability vector (0.0007, 0.0045, 0.0064, 0.9884). If for example we wanted to produce a BAM file with a depth of 3000x, this position would have 2 A, 14 C, 19 G and 2965 T. The probability matrix mentioned in section 2.2.3 is basically the probability vectors of each position of the panel, merged together. In our test and in order to demonstrate our results, we simulated two artificial samples in which we added the calculated background error noise. The first sample or Sample 1 has an average depth of 1000x (+/- 15% at each position) and Sample 2 has an average depth of 10000x. Figure 6 shows how the background error noise is properly and very accurately added at position 2 493 165 of chromosome 1 with the probabilities calculated from the 6 control samples above.

**Fig. 5.**
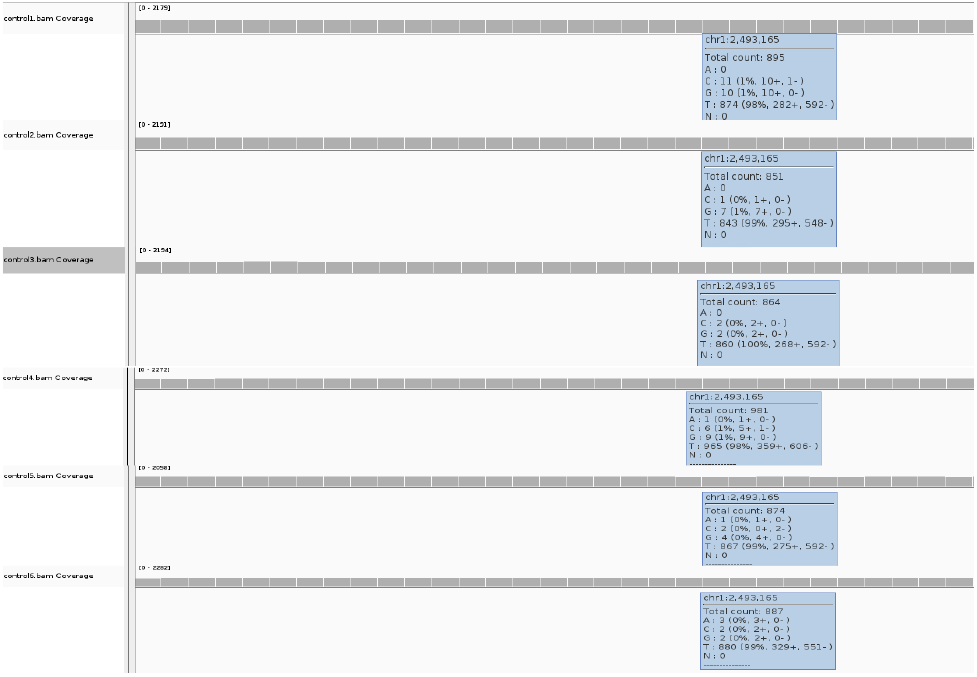
The A,C,G and T breakdown at the position 2 493 165 of the chromosome 1 for the six control samples.

**Fig. 6.**
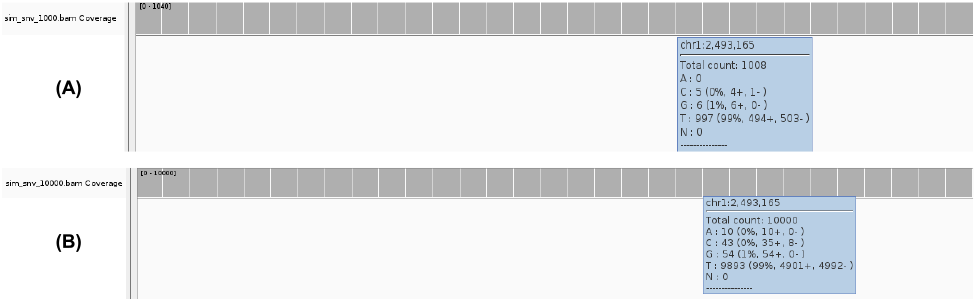
The A, C, G and T breakdown at the position 2 493 165 of the chromosome 1 in the produced samples: Sample 1 with the depth of 1000x (A) and Sample 2 with the depth of10000x (B).

### 3.2 Inserted Variants

Two different lists of mutations were created to go along with each simulated sample. The first list contains 11 substitution variants with frequencies that go from 0.9 (90%) to 0.01 (1%), one deletion at 1% and one insertion at 1%. This list is used to produce the simulated Sample 1 with a depth of 1000x. The second list contains 13 substitution variants with frequencies that go from 0.9 (90%) to 0.001 (0.1%), one deletion at 1% and one insertion at 1%. This list is used to produce the simulated Sample 2 with a depth of 10000x. Two very low frequency variants (frequency < 1%) were added to the second list to test the variant insertion accuracy of UMI-Gen. In fact, very low frequency variants are the hardest to detect and should be systematically used to rigorously test any variant caller. In order to verify that the wanted variants were added at the exact locations with the correct frequencies, we used IGV to visualize the reads. Figure 7 shows and details the variants added in both samples and Table 1 details the exact variants that we inserted at the specific locations. The location of each variant is chosen carefully to make sure that it is not inserted in a long homopolymer region. Figure 8 demonstrates that our tool is capable of accurately adding variants in the final reads at the specified locations for both samples.

**Fig. 7.**
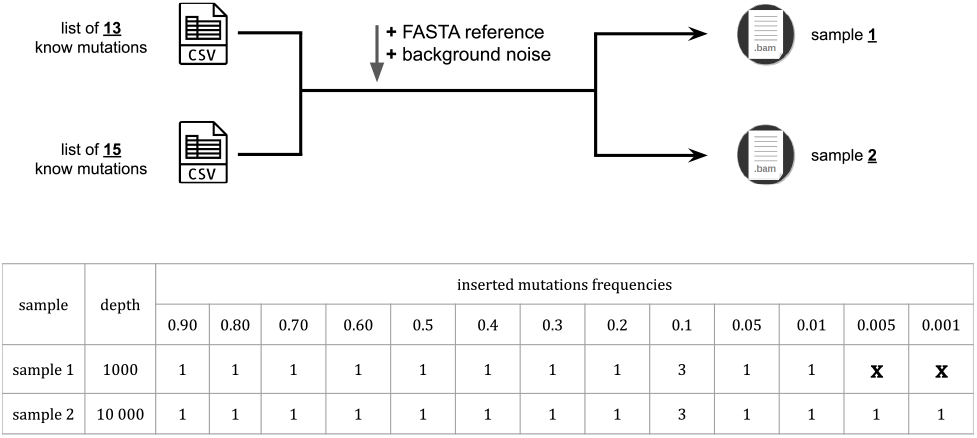
Along with the reference genome FASTA file and the BED file, two different lists were used, one with 13 variants and the other with 15 variants to respectively produce the artificial samples Sample 1 and Sample 2.

**Fig. 8.**
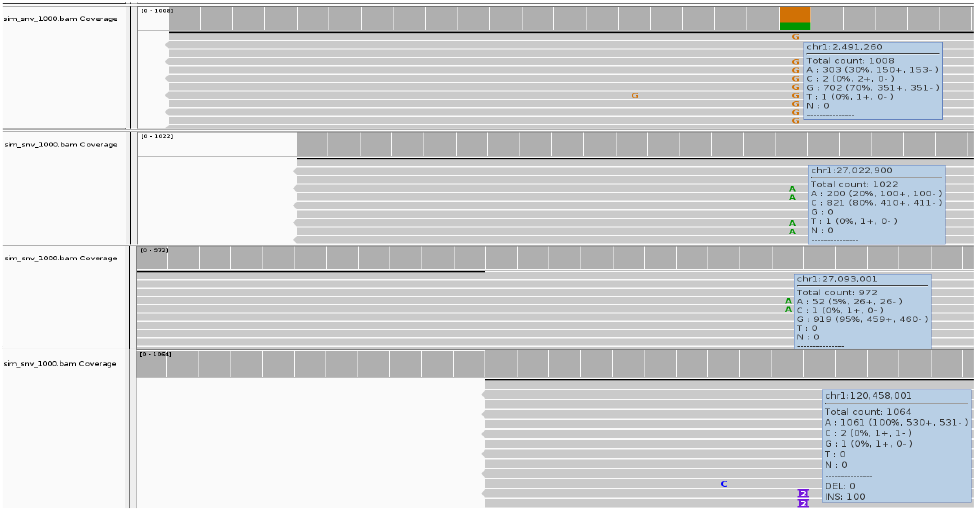
The inserted mutations were correctly added to the reads with their exact locations at their corresponding frequencies. Here, we see four mutations: chr1:2491260A>G at 70%, chr1:27022900C>A at 20%, chr1:120458000C>CTA at 10% and chr1:27093001G>A at 5%.

**Table 1.** Detailed list of the inserted mutations. In this test, all mutations are inserted on chromosome 1.

| Position | Reference allele | Variant allele | Frequency | Sample |
| --- | --- | --- | --- | --- |
| 2 488 101 | G | A | 0.9 | S1 & S2 |
| 2 489 200 | C | A | 0.8 | S1 & S2 |
| 2 491 260 | A | G | 0.7 | S1 & S2 |
| 2 493 201 | T | A | 0.6 | S1 & S2 |
| 2 494 300 | G | A | 0.5 | S1 & S2 |
| 23 885 600 | C | A | 0.4 | S1 & S2 |
| 23 885 800 | A | T | 0.3 | S1 & S2 |
| 27 022 900 | C | A | 0.2 | S1 & S2 |
| 27 023 200 | C | A | 0.1 | S1 & S2 |
| 27 093 001 | G | A | 0.05 | S1 & S2 |
| 27 100 350 | C | A | 0.01 | S1 & S2 |
| 27 106 500 | G | A | 0.005 | S2 only |
| 117 057 400 | T | A | 0.001 | S2 only |
| 120 458 000 | C | CTA | 0.1 | S1 & S2 |
| 120 466 600 | TGTC | T | 0.1 | S1 & S2 |

### 3.3 Variant Detection

We tested the ability of four different variant callers to correctly detect the true variants added in section 3.2 and filter out sequencing errors/artifacts added in 3.1. We used SiNVICT and OutLyzer, two raw-reads-based variant callers specifically developed to detect low frequency variants and two UMI-based variant callers with a very low frequency detection threshold and that analyze UMI tags in order to produce more accurate results. The four variant callers were tested on the two artificial samples: Sample 1 that contains 13 known variants and a depth of 1000x and Sample 2 that contains 15 known variants and a depth of 10000x. Both samples have a total of 76 630 sequenced positions which corresponds to the size of the sequencing panel.

#### 3.3.1 Sample 1

Table 2 details the results of each tool for Sample 1. The total number of positives corresponds to the number of variants found in the result VCF file. The total number of negatives is then calculated by subtracting total positives from the total number of positions (76 630).

**Table 2.** Variant calling results on Sample 1. Four variant callers were tested: SiNVICT, OutLyzer, DeepSNVMiner and UMI-VarCal and for each tool, True Positives (TP), False Positives (FP), False Negatives (FN) and True Negatives (TN) are reported.

| Variant Caller | TP | FP | FN | TN |
| --- | --- | --- | --- | --- |
| SiNVICT | 8 / 241 | 233 / 241 | 5 / 76 389 | 76 384 / 76 389 |
| OutLyzer | 11 / 109 | 98 / 109 | 2 / 76 521 | 76 519 / 76 521 |
| DeepSNVMiner | 0 / 4 | 4 / 4 | 13 / 76 626 | 76 613 / 76 626 |
| UMI-VarCal | 13 / 13 | 0 / 13 | 0 / 76 617 | 76 617 / 76 617 |

Starting with SiNVICT, the tool detected 241 variants in total which corresponds to 76 389 negatives. Among the positives, 8 of them are true positives that can be found in the variants list. However, 233 of the 241 were false positives (96.68 %). Among the negatives, 5 of the 76 389 are false ones. These five variants are in the variants list that we provided and were not detected by the tool. On the other hand, 76 384 positions were correctly not called and considered as true negatives by SiNVICT.

Moving on to OutLyzer, it detected 109 variants in total which corresponds to 76 521 negatives. Among the positives, 11 of them are true positives that can be found in the variants list. However, 98 of the 109 were false positives (89.9 %). Among the 76 521 negatives, only 2 are false ones and 76 519 positions were correctly not called and considered as true negatives by OutLyzer.

Concerning DeepSNVMiner, the tool detected only 4 variants in total which corresponds to 76 626 negatives. Surprisingly, none of the positives are true positives and the 4 positives were all false positives (100 %). Among the negatives, 13 of them are false negatives and 76 613 positions were correctly considered as true negatives by DeepSNVMiner.

Finally, UMI-VarCal was able to detect 13 variants in total which corresponds to 76 617 negatives. All of the detected positives are true positives and no false positives were called by this variant caller (0 %). Among the 76 617 negatives, none is a false negative and 76 617 positions were correctly not called and considered as true negatives by UMI-VarCal.

#### 3.3.2 Sample 2

Table 3 details the results of each tool for Sample 2. Positives and negatives are calculated in the same way as for Sample 1.

**Table 3.** Variant calling results on Sample 2. Four variant callers were tested: SiNVICT, OutLyzer, DeepSNVMiner and UMI-VarCal and for each tool, True Positives (TP), False Positives (FP), False Negatives (FN) and True Negatives (TN) are reported.

| Variant Caller | TP | FP | FN | TN |
| --- | --- | --- | --- | --- |
| SiNVICT | 8 / 463 | 455 / 463 | 7 / 76 167 | 76 160 / 76 167 |
| OutLyzer | 12 / 342 | 330 / 342 | 3 / 76 288 | 76 285 / 76 288 |
| DeepSNVMiner | 0 / 4 | 4 / 4 | 15 / 76 626 | 76 611 / 76 626 |
| UMI-VarCal | 15 / 15 | 0 / 15 | 0 / 76 615 | 76 615 / 76 615 |

Again starting with SiNVICT, the tool detected 463 variants in total which corresponds to 76 167 negatives. Among the positives, 8 of them are true positives that can be found in the variants list. However, 455 of the 463 were false positives (98.27 %). Among the negatives, 7 of the 76 167 are false ones. These seven variants are in the variants list that we provided and were not detected by the tool. On the other hand, 76 160 positions were correctly not called and considered as true negatives by SiNVICT.

Moving on to OutLyzer, it detected 342 variants in total which corresponds to 76 288 negatives. Among the positives, 12 of them are true positives that can be found in the variants list. However, 330 of the 342 were false positives (96.49 %). Among the 76 288 negatives, only 3 are false ones and 76285 positions were correctly not called and considered as true negatives by OutLyzer.

Concerning DeepSNVMiner, the tool detected only 4 variants in total which corresponds to 76 626 negatives. Surprisingly again, we obtained the same 4 false positives from Sample 1 with no true positives. Among the negatives, 15 of them are false negatives and 76 611 positions were correctly considered as true negatives by DeepSNVMiner.

Finally, UMI-VarCal was able to detect 15 variants in total which corresponds to 76 615 negatives. Once again, all of the detected positives are true positives and no false positives were called by the tool (0 %). Among the 76 615 negatives, none is a false negative meaning that all of them were correctly not called and considered as true negatives by UMI-VarCal.

## 4 Discussion

Tagging DNA fragments with UMI tags have proved itself as a very reliable method to significantly reduce - if not completely remove - the number of false positives upon variant calling. A huge number of variant callers are publicly available at the moment but unfortunately, only 4 of them are specifically developed to treat UMI tags in reads. For raw-reads-based variant callers, a lot of artificial reads simulator exist and can satisfy everyone’s needs. However, at our knowledge, no tool is publicly available to simulate artificial reads with UMI tags. Such tool is very important as it allows developers to accurately test the specificity and sensitivity of their variant callers on artificial reads in which real variants are known instead of testing them on biological samples whose mutational profile is completely or partially unknown.

Our main objective was to develop a UMI-tagged reads simulator that is fast, accurate and reliable. UMI-Gen is able to estimate the background error noise of a given control dataset and then reproduce it accurately in the produced reads. Doing so, it allows to mimic the sequencer’s background noise of a real sequencing experiment. We also showed that our simulator is able to accurately insert variants if provided with a list of variants with exact locations and their corresponding frequencies. In our tests, we were able to insert mutations as low as 0.1% but theoretically, we can go as low as we want provided that the depth of the produced sample is accordingly increased.

Moreover, in our variant callers comparison, SiNVICT did a somewhat good job detecting the 8 of the added variants and went as low as 5%. Impressively, we judge the performance of OutLyzer as excellent as it detected 12 of the 15 variants (Sample 2) and showed a detection threshold of 0.5% which is very respectable. However, SiNVICT and OutLyzer being raw-reads-based variant callers, UMI tags were not treated in the reads and therefore, both tools produced a very high percentage of false positives. DeepSNVMiner results were surprising since it couldn’t detect any inserted variants and only called out 4 false positives. This is rather weird given that it is supposed to be a variant caller that treats UMI tags and should show increased sensitivity and specificity when compared to raw-read-based ones. UMI-VarCal on the other hand was successfully able to treat UMI tags allowing it to filter out all false positives and only call out the 13 added variants in Sample 1 and all of the 15 in Sample 2. These results demonstrate UMI-VarCal’s very high sensitivity and specificity on both samples. It performed better than the three tools we tested and showed a detection threshold of 0.1%.

## 5 Conclusion

Here, we present UMI-Gen: a standalone UMI-based reads simulator for variant calling evaluation in paired-end sequencing NGS libraries. UMI-Gen produces sequencing files (FASTQ, BAM and SAM) for an artificial sample to be used for UMI-based variant calling testing purposes. By using a set of control DNA samples, our tool is capable to accurately mimic the background error noise of the sequencer and add it into the reads. After that, it can insert specific mutations at specific locations and at very precise frequencies that can go as low as 0.1% (and even lower). In our tests, all added artifacts were correctly inserted in the reads, causing a high number of false positives in the raw-reads-based variant callers results. Also, all inserted true variants were visualized with a genome visualizing tool (IGV) and were detected by at least one of the four variant calling tools we tested. Finally, we note that UMI-Gen’s filters and parameters (such as read length and UMI tag length) are customizable which gives the user total control over his produced samples. This level of customization allows the tool to be adequate for a high number of research applications.

## References

Andrews, T. D., Jeelall, Y., Talaulikar, D., Goodnow, C. C., and Field, M. A. (2016). DeepSNVMiner: a sequence analysis tool to detect emergent, rare mutations in subsets of cell populations. PeerJ, 4.

Bar, D. Z., Arlt, M. F., Brazier, J. F., Norris, W. E., Campbell, S. E., Chines, P., Larrieu, D., Jackson, S. P., Collins, F. S., Glover, T. W., and Gordon, L. B. (2017). A novel somatic mutation achieves partial rescue in a child with Hutchinson-Gilford progeria syndrome. J Med Genet, 54(3), 212–216.

Kockan, C., Hach, F., Sarrafi, I., Bell, R. H., McConeghy, B., Beja, K., Haegert, A., Wyatt, A. W., Volik, S. V., Chi, K. N., Collins, C. C., and Sahinalp, S. C. (2017). SiNVICT: ultra-sensitive detection of single nucleotide variants and indels in circulating tumour DNA. Bioinformatics, 33(1), 26–34.

Kukita, Y., Matoba, R., Uchida, J., Hamakawa, T., Doki, Y., Imamura, F., and Kato, K. (2015). High-fidelity target sequencing of individual molecules identified using barcode sequences: de novo detection and absolute quantitation of mutations in plasma cell-free DNA from cancer patients. DNA Res, 22(4), 269–277.

Li, H. and Durbin, R. (2009). Fast and accurate short read alignment with burrows-wheeler transform. Bioinformatics, 25(14), 1754–1760.

Li, H., Handsaker, B., Wysoker, A., Fennell, T., Ruan, J., Homer, N., Marth, G., Abecasis, G., and Durbin, R. (2009). The Sequence Alignment/Map format and SAMtools. Bioinformatics, 25(16), 2078–2079.

Muller, E., Goardon, N., Brault, B., Rousselin, A., Paimparay, G., Legros, A., Fouillet, R., Bruet, O., Tranchant, A., Domin, F., San, C., Quesnelle, C., Frebourg, T., Ricou, A., Krieger, S., Vaur, D., and Castera, L. (2016). OutLyzer: software for extracting low-allele-frequency tumor mutations from sequencing background noise in clinical practice. Oncotarget, 7(48), 79485–79493.

Newman, A. M., Lovejoy, A. F., Klass, D. M., Kurtz, D. M., Chabon, J. J., Scherer, F., Stehr, H., Liu, C. L., Bratman, S. V., Say, C., Zhou, L., Carter, J. N., West, R. B., Sledge, G. W., Shrager, J. B., Loo, B. W., Neal, J. W., Wakelee, H. A., Diehn, M., and Alizadeh, A. A. (2016). Integrated digital error suppression for improved detection of circulating tumor DNA. Nat Biotechnol, 34(5), 547–555.

Robinson, J. T., Thorvaldsdóttir, H., Winckler, W., Guttman, M., Lander, E. S., Getz, G., and Mesirov, J. P. (2011). Integrative Genomics Viewer. Nature biotechnology, 29(1), 24–26.

Sater, V., Viailly, P.-J., Lecroq, T., Prieur-Gaston, E., Bohers, E., Viennot, M., Ruminy, P., Dauchel, H., Vera, P., and Jardin, F. (2020). UMI-VarCal: a new UMI-based variant caller that efficiently improves low-frequency variant detection in paired-end sequencing NGS libraries. Bioinformatics (Oxford, England).

Schmitt, M. W., Kennedy, S. R., Salk, J. J., Fox, E. J., Hiatt, J. B., and Loeb, L. A. (2012). Detection of ultra-rare mutations by next-generation sequencing. Proc Natl Acad Sci U S A, 109(36), 14508–14513.

Young, A. L., Challen, G. A., Birmann, B. M., and Druley, T. E. (2016). Clonal haematopoiesis harbouring AML-associated mutations is ubiquitous in healthy adults. Nat Commun, 7.

