## supplementary for "UMI-Gen: a UMI-based reads simulator for variant calling evaluation in paired-end sequencing NGS libraries"

### 1 Tables

| **Symbol** | **ASCII Code** | **Q-Score** | **Symbol** | **ASCII Code** | **Q-Score** |
|---|---|---|---|---|---|
| ! | 33 | 0 | 6 | 54 | 21 |
| “ | 34 | 1 | 7 | 55 | 22 |
| # | 35 | 2 | 8 | 56 | 23 |
| $ | 36 | 3 | 9 | 57 | 24 |
| % | 37 | 4 | : | 58 | 25 |
| & | 38 | 5 | ; | 59 | 26 |
| ’ | 39 | 6 | < | 60 | 27 |
| ( | 40 | 7 | = | 61 | 28 |
| ) | 41 | 8 | > | 62 | 29 |
| * | 42 | 9 | ? | 63 | 30 |
| + | 43 | 10 | @ | 64 | 31 |
| , | 44 | 11 | A | 65 | 32 |
| - | 45 | 12 | B | 66 | 33 |
| . | 46 | 13 | C | 67 | 34 |
| / | 47 | 14 | D | 68 | 35 |
| 0 | 48 | 15 | E | 69 | 36 |
| 1 | 49 | 16 | F | 70 | 37 |
| 2 | 50 | 17 | G | 71 | 38 |
| 3 | 51 | 18 | H | 72 | 39 |
| 4 | 52 | 19 | I | 73 | 40 |
| 5 | 53 | 20 | J | 74 | 41 |

Table S1: Quality scores encoding

| **Parameter** | **Type** | **Required** | **Default** |
|---|---|---|---|
| --input (-i) | list of files | YES | NONE |
| --fasta (-f) | file | YES | NONE |
| --bed (-b) | file | YES | NONE |
| --pileup (-p) | file | | NONE |
| --variants (-v) | file | | NONE |
| --output (-o) | dir | | ./ |
| --name (-n) | string | | sim_$< depth_value >$ |
| --min_base_quality | int | | 10 |
| --min_read_quality | int | | 20 |
| --min_mapping_quality | int | | 20 |
| --min_variant_umi | int | | 10 |
| --alpha | float | | 0.05 |
| --strand_bias_method | string | | default |
| --max_strand_bias | float | | 1 |
| --max_hp_length | int | | 7 |
| --depth (-d) | int | | 1000 |
| --umi_length (-u) | int | | 12 |
| --read_length | int | | 110 |
| --max_noise_rate | float | | 0.05 |
| --amp_factor | int | | 10 |

Table S2: List of all UMI-Gen parameters. A detailed explication of each parameter is found in the tool documentation.

| **Gene** | **Number of regions** | **Gene** | **Number of regions** |
|---|---|---|---|
| ARID1A | 85 | GNA13 | 14 |
| B2M | 6 | ID3 | 6 |
| BCL2 | 8 | IRF4 | 22 |
| BRAF | 2 | MEF2B | 21 |
| BTK | 3 | MYC | 16 |
| CARD11 | 16 | MYD88 | 8 |
| CCND3 | 15 | NOTCH1 | 19 |
| CD58 | 17 | NOTCH2 | 24 |
| CD79A | 5 | PIM1 | 14 |
| CD79B | 4 | PLCG2 | 16 |
| CDKN2A | 27 | PRDM1 | 33 |
| CDKN2B | 47 | SOCS1 | 8 |
| CIITA | 66 | STAT6 | 14 |
| CREBBP | 103 | TCF3 | 5 |
| CXCR4 | 12 | TNFAIP3 | 31 |
| EP300 | 102 | TNFRSF14 | 16 |
| EZH2 | 5 | TP53 | 24 |
| FOXP1 | 22 | XPO1 | 11 |

Table S3: Pan-lymphoma Panel: List of targeted regions per gene
